# The winner takes it all: how semelparous insects can become periodical

**DOI:** 10.1101/446252

**Authors:** Odo Diekmann, Robert Planqué

## Abstract

The aim of this short note is to give a simple explanation for the remarkable periodicity of *Magicicada* species, which appear as adults only every 13 or 17 years, depending on the region. We show that a combination of two types of density dependence may drive, for large classes of initial conditions, all but one year class to extinction. Competition for food leads to negative density dependence in the form of a uniform (i.e., affecting all age classes in the same way) reduction of the survival probability. Satiation of predators leads to positive density dependence within the reproducing age class. The analysis focuses on the full life cycle map derived by iteration of a semelparous Leslie matrix.

## 1 Periodical insects: a conundrum

According to M. G. Bulmer (Bulmer 1977), “An insect population is said to be periodical if the life cycle has a fixed length of *k* years (k > 1) and if the adults do not appear every year but only every kth year.” He then provides several examples, one by quoting Lloyd and Dybas (1966)

> “Periodical cicadas, *Magicicada* spp., have the longest life cycles known for insects. In any one population, all but a tiny fraction are the same age. The nymphs suck juices from the roots of deciduous forest trees in eastern United States. Mature nymphs finally emerge from the ground, become adults, mate, lay their eggs, and die within the same few weeks of every seventeenth (or, in the South, every thirteenth) year. Not one species does this, but three. There are three separate and distinct species that occur together over most of the range of periodical cicadas and, wherever the species coexist, they are invariably synchronized. In different regions, different “broods” of periodical cicadas may be out of synchrony by several years, but the species (in a given region) never are. Finally, the same three species—the same as nearly as anyone can tell by looking at them or listening to the songs of their males—exist both as 17 and as 13-year periodical cicadas! [Pp. 133 - 134; Note that the three species are now considered to be seven species, each with their own period of either 13 or 17 years (Marshall 2008)]”

In the wake of Bulmer’s pioneering paper a rich literature on discrete time models for semelparous organisms developed, see (Behncke 2000, Cushing 2009; 2015, Cushing and Henson 2012, Diekmann et al. 2005, Kon 2012; 2017, Webb 2001, Wikan 2017) and the references given there. The population splits into year classes according to the year of birth (equivalently: the year of reproduction) counted modulo *k*. As a year class is reproductively isolated from the other year classes, it forms a population by itself. Accordingly, competition may lead to exclusion.

Despite this rich literature, the spatio-temporal pattern displayed by periodical cicada population dynamics is, we think, still an enigma. The interest of one of the authors in the subject was rekindled while refereeing an early version of (Machta et al. 2019). The model introduced and studied in that paper is characterized by

– uniform (i.e., age-class unspecific) negative density dependence during development;
– positive density dependence for reproduction.

The first of these captures competition for food, the second satiation of predators to eat adults in a given year (Williams et al. 1993).

The paper (Machta et al. 2019) considers the limit *k* → ∞ and employs a continuous time description of the competition for resources (whence the word “hybrid” in the title). The main result establishes the instability of any steady state with more than one year class present.

Here we focus on the “full life cycle” map, i.e., the k-times iterated Leslie matrix, cf. (Davydova et al. 2003; 2005). The full life cycle map is represented by a diagonal matrix. If competition is uniform, in the sense that in any year any of the survival probabilities is the product of a fixed age-class specific factor and a year-specific factor that is the same for all age-classes, the diagonal elements all have the same overall survival factor. In other words, negative density dependence results over a full life cycle for all year classes in exactly the same multiplication factor. Each diagonal element has an additional reproduction factor. Positive density dependence causes this factor to be highest for the year class that had, in the year that it reproduced, the highest density. In this way, a numerical advantage is reinforced and asymptotically there remains, “generically”, only one year class: all the others go extinct. See Figure 1 for an example.

**Fig 1.**
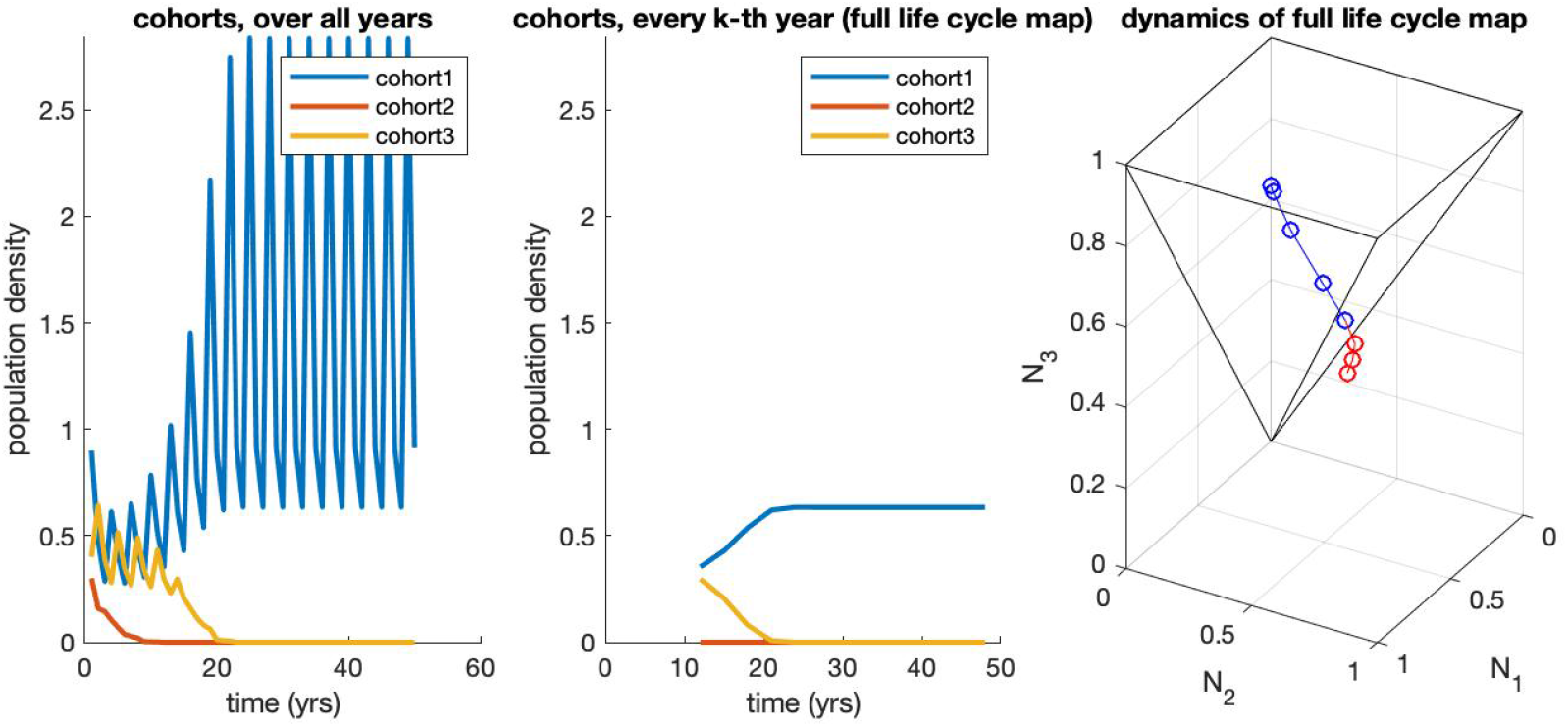
Example dynamics for the map (1)–(2), with parameters *σ*_1_ = *σ*_2_ = *σ*_3_ = 1, *π*(***N***) = *e*^−0.4(*N*_1_+*N*_2_+*N*_3_)^ and *β*(*N*_3_) = 17.5*N*_3_/(1 + *N*_3_). Left: full dynamics of all cohorts; middle: the dynamics of the full life cycle map, for all cohorts, shown from the year at which the orbit enters *Ω*_3_; right: dynamics generated by the full life cycle map, with *Ω*_3_ in black, with red points of the orbit outside *Ω*_3_ and blue those inside. We observe convergence to a single year class fixed point inside *Ω*_3_ that lies on the *N*_3_-axis. Note that the choice *σ*_1_ = *σ*_2_ = *σ*_3_ = 1 is no restriction, as we can scale the variables *N_k_* with these parameters, as indicated in Section 4. Do note, however, that stronger mortality increases the size of *Ω_k_*.

Note that this holds independently of the resulting single year class (SYC) dynamics in the sense that the winners may exhibit ultimately steady state, periodic or even chaotic dynamics. Also note that it depends on the initial condition which one of the *k* year classes will win the competition. Domains of attraction (or, dominance) are separated by sets of special initial conditions for which several year classes coexist forever (e.g., steady coexistence states and their stable manifolds). In principle these separatrices may have an intricate structure (e.g., see (Hofbauer et al. 2004)).

The upshot is that the combination of uniform negative density dependence and concentrated (in one point of the life cycle) positive density dependence, as assumed in (Machta et al. 2019), leads, as a rule, to exclusion of all but at most one year class. This does not prove, of course, that this is the mechanism underlying the observed phenomenon of SYC dynamics in *Magicicada*. But in the spirit of Occam’s Razor, this probably currently offers the simplest, and hence most plausible, potential explanation.

Anyhow, as far as we are aware, we provide below the first analytical demonstration of one year class driving all other year classes to extinction without severe restrictions on the initial conditions and without any condition on the resulting SYC dynamics, except for boundedness.

## 2 Model formulation

Let *k* be an integer bigger than one. Let ***N***(*t*) be the *k*-vector such that *N_j_*(*t*) is the sub-population density of individuals that at the census moment in year *t* have age *j*. In view of linear algebra we take *j* = 1, 2,…, *k*, even though biologically *j* = 0,1,…, *k* – 1 would perhaps make more sense if, as we assume, the census moment is in the early autumn, so after reproduction took place. We assume that

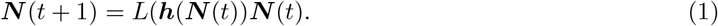

where

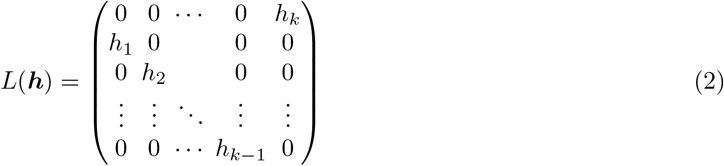

is a “semelparous” Leslie matrix (here the adjective “semelparous” indicates that in the first row only the last element is non-zero, reflecting that individuals reproduce at age *k* and then die). The dependence of the matrix elements *h_i_* on ***N*** captures density dependence. We assume that

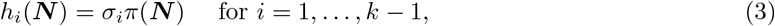

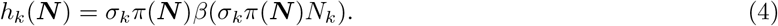

where

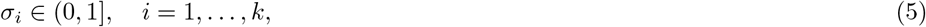

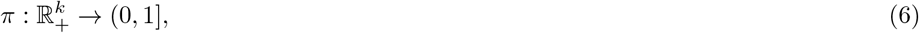

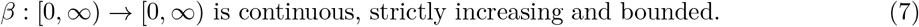

Here the *σ_i_* are survival probabilities under “standard” conditions and the (uniform, i.e., *i*-independent) factor *π*(***N***) reflects the reduction of the survival probability as a result of crowding and competition for food. The last age group too needs to survive the winter before they can emerge as adults, so the population density of emerging adults equals *σ_k_π*(***N***)*N_k_*. The function *β* is a composite model ingredient. It describes the number of offspring at the census moment per emerging adult. The monotonicity of *β* reflects that the per adult capita risk of falling victim to predation decreases with adult population density. This is the effect of predators becoming satiated (or even over-satiated) when prey density is very high, and that the time interval between emergence and death of the adults is too short for a numerical increase in the number of predators (mainly birds).

## 3 The main results

So far we did not specify any property of *π* that justifies the description “negative” density dependence. But now we require that population densities remain bounded and implicitly this is an assumption concerning the function *π*.

### Hypothesis 1

*(dissipativity (Hale 1988)) There exists R* > 0 *such that*

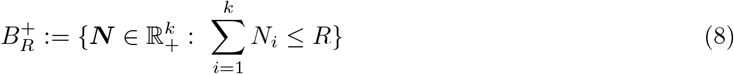

*is forward invariant and attracts all orbits*.

By “attracts all orbits” we mean that for every 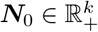 there exists *m* = *m*(***N***_0_) such that the solution of (1) with initial condition ***N***(0) = ***N***_0_ satisfies 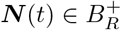 for *t* ≥ *m*(***N***_0_). We adopt Hypothesis 1 for the rest of the paper. In Section 4 below we provide easily verifiable conditions on *π* and *β* that guarantee that Hypotheses 1 is satisfied.

At the end of Section 1 we already observed that (due to symmetry, see (Davydova et al. 2005, Diekmann and van Gils 2003) for a bit more detail) any year class is a candidate for becoming the sole inhabiter of the world. It turns out to be relatively easy to describe a set of initial conditions such that the year class that has age *k* at the initial time will outcompete all the other year classes.

The underlying idea is the following. During a full life cycle, a year class has highest density at the first census after birth and lowest density at the last census before reproduction, simply since inbetween the density is reduced by mortality. Likewise, for a steady state with all year classes present, abundance decreases with age. (Possibly the same holds for any solution of period *k* with several year classes present, but we do not know.) The age classes at a particular census, on the other hand, reflect the relative abundances of the year classes, albeit in a manner that does not allow general straightforward meaningful comparison. However, if the highest age class has highest abundance, this clearly shows that the corresponding year class is dominant and on the way to becoming the winner. As shown in the following theorem, this test can be refined by correcting for the predictable mortality (as captured by the *σ*’s) before reaching the highest age.

### Theorem 1

*The set*

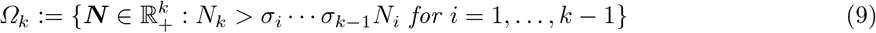

*is forward invariant under the full life cycle map, and if **N***(0) ∈ *Ω_k_ and for given i* ∈ {1,…, *k* – 1}, *N_i_*(0) > 0, *then*

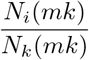

*is a strictly decreasing function of m. This function has limit zero for m* → ∞ *in case* limsup_*m*→∞_ *N_k_*(*mk*) > 0.

The set *Ω*_3_ is depicted in Figure 1. Note that *all* year classes go extinct if *N_k_*(*mk*) → 0 for *m* → ∞. Hypothesis 1 rules out the possibility that both *N_i_*(*mk*) and *N_k_*(*mk*) grow beyond any bound for *m* → ∞ while their ratio goes to zero. So we conclude that for initial conditions belonging to *Ω_k_* only the *k*-th year class can possibly persist.

### Proof

In order to avoid overburdening the reader with notational detail, we focus on the proof of the case *k* = 3. The proof for general *k* does not require new arguments.

From (1) it follows that

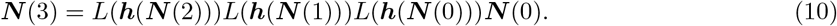

The product of the three Leslie matrices is a diagonal matrix. We claim that the three elements of the diagonal of this matrix have a factor

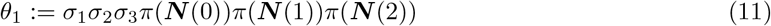

in common and are in fact of the form *β*(·)*θ*_1_ with the argument of *β* being *θ*_1_*N*_1_(0) for the first element, *θ*_2_*N*_2_(0) with *θ*_2_:= *σ*_2_*σ*_3_*π*(***N***(0))*π*(***N***(1)) for the second element, and *θ*_3_*N*_3_(0) with *θ*_3_:= *σ*_3_*π*(***N***(0)) for the third element.

To verify the claim, we elaborate (10) from right to left, using *π*_1_ to denote *π*(***N***(0)), *π*_2_ to denote *π*(***N***(1)) and *π*_3_ to denote *π*(***N***(2)). All factors in the products in the vectors below follow the order of events over time.

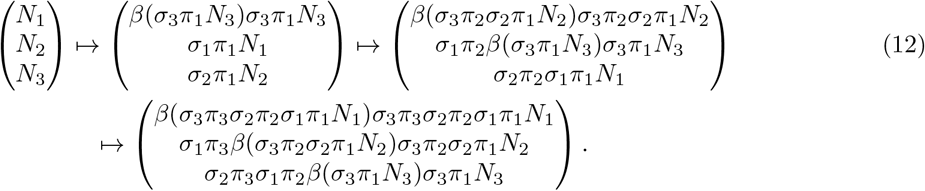

Noting that *θ*_1_ = *σ*_1_*σ*_2_*σ*_3_*π*_1_*π*_2_*π*_3_, inspection reveals that the claim is correct. It follows that

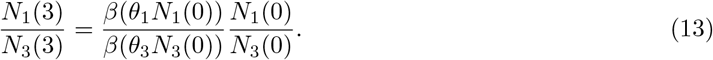

Now for ***N***(0) ∈ *Ω*_3_, *σ*_1_*σ*_2_*N*_1_(0) < *N*_3_(0), so that

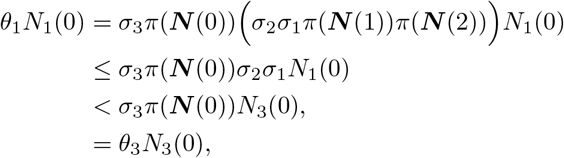

and we conclude that, since *β* is strictly increasing,

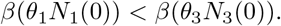

Hence, when *N*_1_(0) > 0,

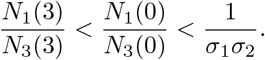

Essentially the same argumentation yields that, when ***N***(0) ∈ *Ω*_3_ and *N*_2_(0) > 0, then

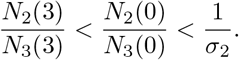

It follows that *Ω*_3_ is forward invariant under the full life cycle map and that for *i* = 1, 2 the sequences

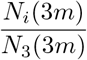

are decreasing in *m*, strictly when positive. It remains to be shown that either their limit is zero or all year classes go extinct.

Assume that *N*_1_(3*m*)/*N*_3_(3*m*) → *α* for *m* → ∞. Extending the definitions of *θ_i_* to

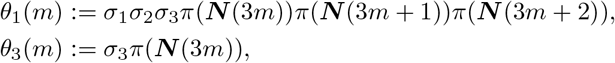

then by iteration (13) leads to

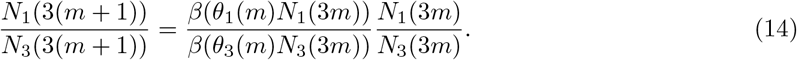

Passing to the limit *m* → ∞ in (14), it follows that if *α* > 0 then necessarily, since *β* is continuous,

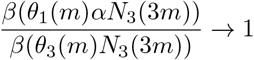

for *m* → ∞. This is very well possible if *N*_3_(3*m*) → 0 for *m* →∞ (and in that case, since *Ω*_3_ is forward invariant, also *N*_1_(3*m*) and *N*_2_(3*m*) converge to zero for *m* → ∞, so all year classes go extinct). But if limsup_*m*→∞_ *N*_3_(3*m*) > 0, this is *not* possible, since *α* < 1, *θ*_1_(*m*) ≤ *θ*_3_(*m*) because *σ*_1_*σ*_2_ ≤ 1, *π*(***N***(3*m* + 1))*π*(***N***(3*m* + 2)) ≤ 1, and *β* is strictly increasing.

Theorem 1 does neither restrict nor predict the resulting SYC dynamics. But does it allow for a large class of initial conditions? The answer, of course, heavily depends on what one means by “large”. Certainly the set is open. Moreover, if a given initial condition does not belong to *Ω_k_*, one can apply (1) once and check whether ***N***(1) belongs to *Ω_k_* (if it does, the year class that had label *k* – 1 in year 0 is the winner). By applying (1) repeatedly one can thus check for any of the year classes whether or not Theorem 1 guarantees that they will win the competition. The example of a steady state with all year classes present clearly shows that this procedure may fail to point out a winner, for the simple reason that there might not be a winner. The next question then is: how exceptional is it that several year classes persist?

It is tempting to conjecture that a fixed point of the full life cycle map, with more than one year class having positive density, is necessarily unstable, as was found to be the case for the model considered by Machta et al. (2019). However, in the Appendix we show that such a fixed point can in fact be stable if the negative density dependence incorporated in *π* is stronger than the positive density dependence incorporated in *β*, in the sense that an *increase* of the oldest age group *N_k_* can lead to a decrease of *π*(***N***)*N_k_* and thus to a *decrease* of the youngest age group in the next year. So without further restrictions on the class of models, it is *not* guaranteed that generically there will be a single winner. The restriction that for all non-negative *N*_1_,…, *N*_*k*–1_ the map *N_k_* ↦ *π*(***N***)*N_k_* is increasing seems both meaningful and promising. See the Appendix for an example.

As illustrated in Figure 2, numerical tests indicate that the occurrence of a single winner is in fact rather common. In the same figure we also evaluate the performance of the following

**Fig 2.**
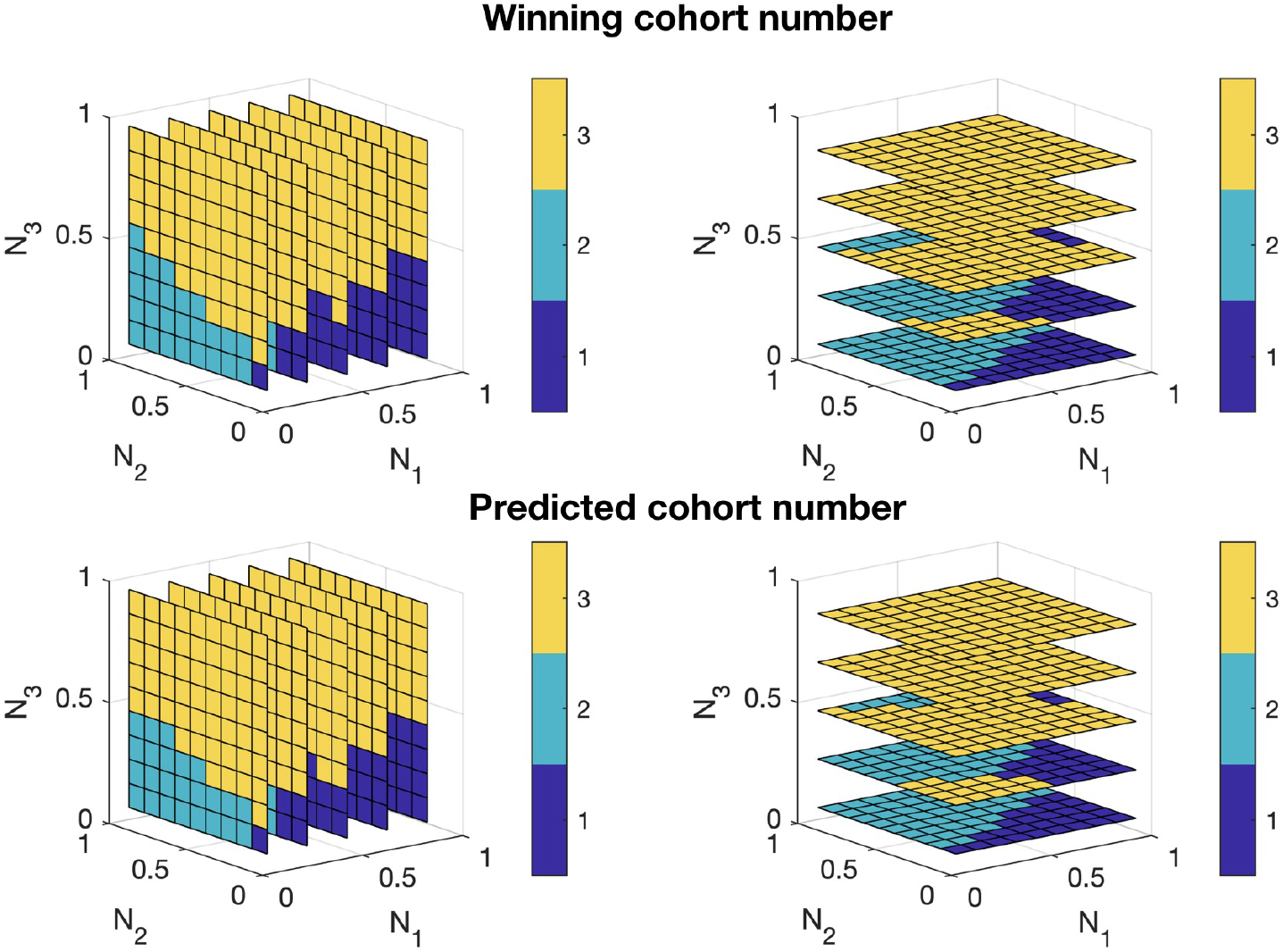
The long-term dynamics as determined numerically, illustrating that for almost all initial conditions, a single year class emerges as the “winner”. Top row: the winning cohort number, as a function of initial values (*N*_1_(0), *N*_2_ (0), *N*_3_(0)), shown in slices in the *N*_1_-direction (left) and in the *N*_3_-direction (right). Bottom row: the cohort number that is predicted to win using the Prediction Tool in the text. This tool is good, but not perfect, in predicting which cohort will eventually rule the world. Model ingredients are *β*(*x*) = max{0, 10(1 – 0.1/*x*)}, and *π*(***N***) = e^−0.7(*N*_1_+*N*_2_+*N*_3_)^, and *σ*_1_ = *σ*_2_ = *σ*_3_ = 1.

### Prediction Tool

*For an arbitrary initial condition **N***(0), *find the index j that maximizes the diagonal elements in the full life cycle map, i.e*.,

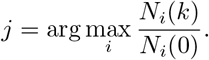

*The prediction is that the year class having age j in year 0 will drive all other year classes to extinction*.

It appears that the tool works well, but is not infallible.

We now present a theorem in which we incorporate more restrictions, but also describe more precisely the ultimate dynamics. The full life cycle map acts on ℝ^*k*^. Any subspace characterised by *k*–1 components being zero is forward invariant. The restriction to such a forward invariant subspace corresponds to a map from ℝ to ℝ that we shall call a SYC full life cycle map. The *k* maps are different but equivalent, see (Davydova 2004, Chapter V; Davydova et al. 2003, Section 7). To stay in line with Theorem 1 we work primarily with the map corresponding to *N*_1_ = *N*_2_ = … = *N*_*k*–1_ = 0.

#### Theorem 2

*If a single year class fixed point or periodic orbit of the SYC full life cycle map is linearly stable with respect to that map, so with respect to within year class perturbations, it is automatically also linearly stable with respect to perturbations that involve small introductions of other year classes*.

*Proof* We again give a proof for the case *k* = 3. The proof for general *k* uses no new arguments.

The full life cycle map given by (12) is represented by a diagonal 3 × 3 matrix *M* acting on a 3 vector,

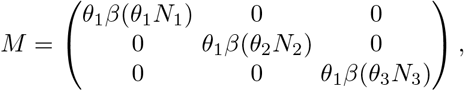

where, as before,

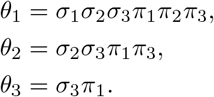

(Of course, all the *π_i_* are functions of *N*_1_, *N*_2_, and *N*_3_.) We first consider a SYC fixed point and choose the phase such that only the third component of the fixed point vector is positive. Then necessarily *M*_3,3_ = *θ*_1_*β*(*θ*_3_*N*_3_) = 1 when evaluated in the fixed point. *N*_3_ is itself a fixed point of the SYC map

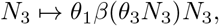

and is, by assumption, stable as such.

The linearisation of the full life cycle map in the fixed point is represented by a matrix as well, of course, *L*, say. Since the first two components of the fixed point are zero, *L*_1,2_ = *L*_1,3_ = *L*_2,3_ = *L*_2,1_ = 0, making *L* a lower-diagonal matrix, with eigenvalues equal to the diagonal elements. Direct computation shows that *L* must be of the form

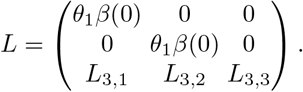

Since *β* is increasing, 0 ≤ *θ*_1_*β*(0) < *θ*_1_ *β*(*θ*_3_*N*_3_) = *M*_3,3_ = 1.

The diagonal element in the third and last row is complicated, but is equal to the linearisation of the SYC full life cycle map in the fixed point. Hence, since we have assumed that fixed point to be linearly stable, |*L*_3,3_| < 1. So all eigenvalues of *L* are less than one in absolute value and the fixed point of the three dimensional full life cycle map is linearly stable.

For an orbit of period p, we consider the *p* times iterated full life cycle map. The structure is exactly the same as before, the only difference is that diagonal elements have *p* factors *θ*_1_*β*. The Jacobi matrix is again lower-diagonal, and of the form

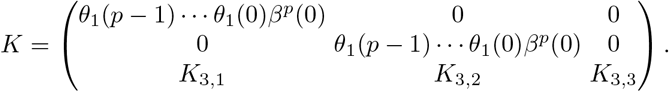

By the same arguments as before, *θ*_1_(*p* – 1) · · · *θ*_1_(0)*β^p^*(0) < 1, and |*K*_3,3_| < 1 by the assumption that the *p*-periodic orbit is linearly stable. So exactly as before we reach the conclusion that all eigenvalues are less than one in absolute value.

## 4 Model ingredients

To obtain more insight which choices of model ingredients *π* and *β* ensure persistence of populations and induce dissipative dynamics as defined in Hypothesis 1, we scale the variables in the following way. Let

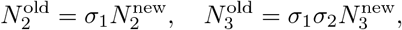

and define functions *β*^new^ and *π*^new^ according to

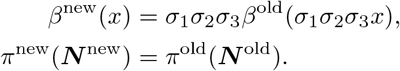

In terms of these new variables, and dropping the superscripts for convenience, the map is seen to be simplified to

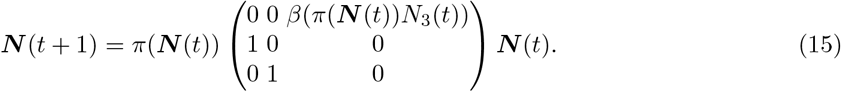

Let us now choose the following form for *π*:

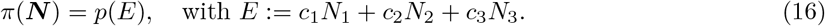

So *π*(***N***) is determined by first computing a weighted population size *E* and next applying a scalar map *p* that assigns to *E* a component of the survival probability, i.e., a number between zero and one. Assume *c*_3_ > 0, and define

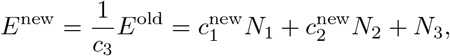

and

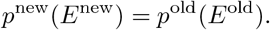

Let us now consider the SYC dynamics. To facilitate the description of that case, we introduce

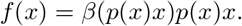

Dropping again the “new” superscripts, the SYC dynamics are given by

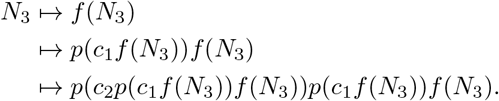

To ensure boundedness of the population consisting of just one year class, the graph of the SYC full life cycle map,

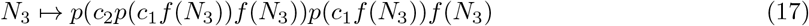

should be below the 45° line for large values of *N*_3_. Our model ingredients *β* and *p* must be chosen accordingly. Figure 3 gives an impression of the form of the graph of the SYC full life cycle map, for different choices of the model ingredients.

**Fig 3.**
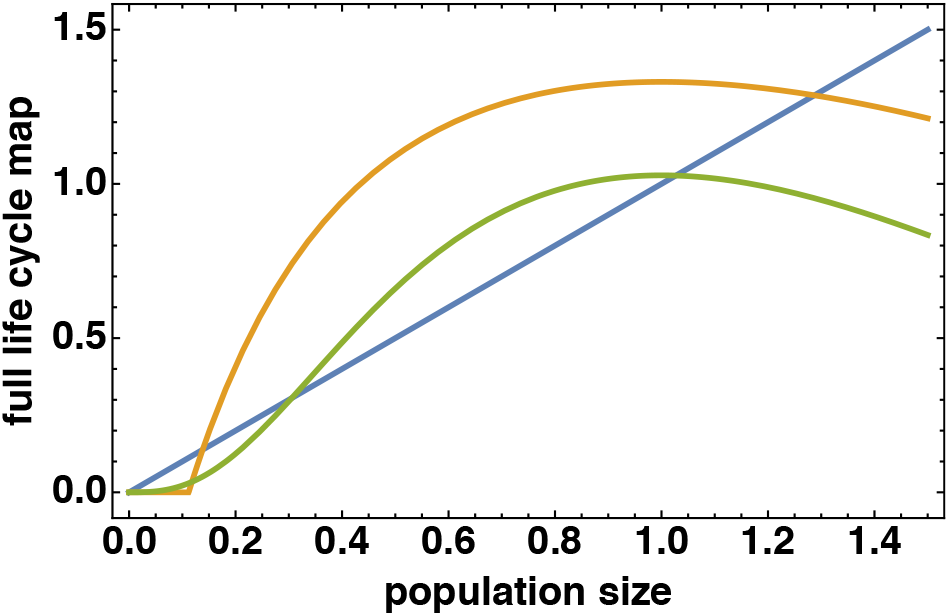
Example graphs of the SYC full life cycle map (17) for different choices of model ingredients. The blue line shows the diagonal; for both full life cycle map graphs, *π*(*E*) = *e*^∞*E*^; the orange curve uses *β*(*x*) = max{0, 7(1 – 0.1/*x*)}, the green curve *β*(*x*) = 30*x*^2^/(1 + *x*^2^). Further parameters chosen in (17) are *c*_1_ = *c*_2_ = 0.1, *c*_3_ = 1. Note that both maps show an Allee effect.

**Fig 4.**
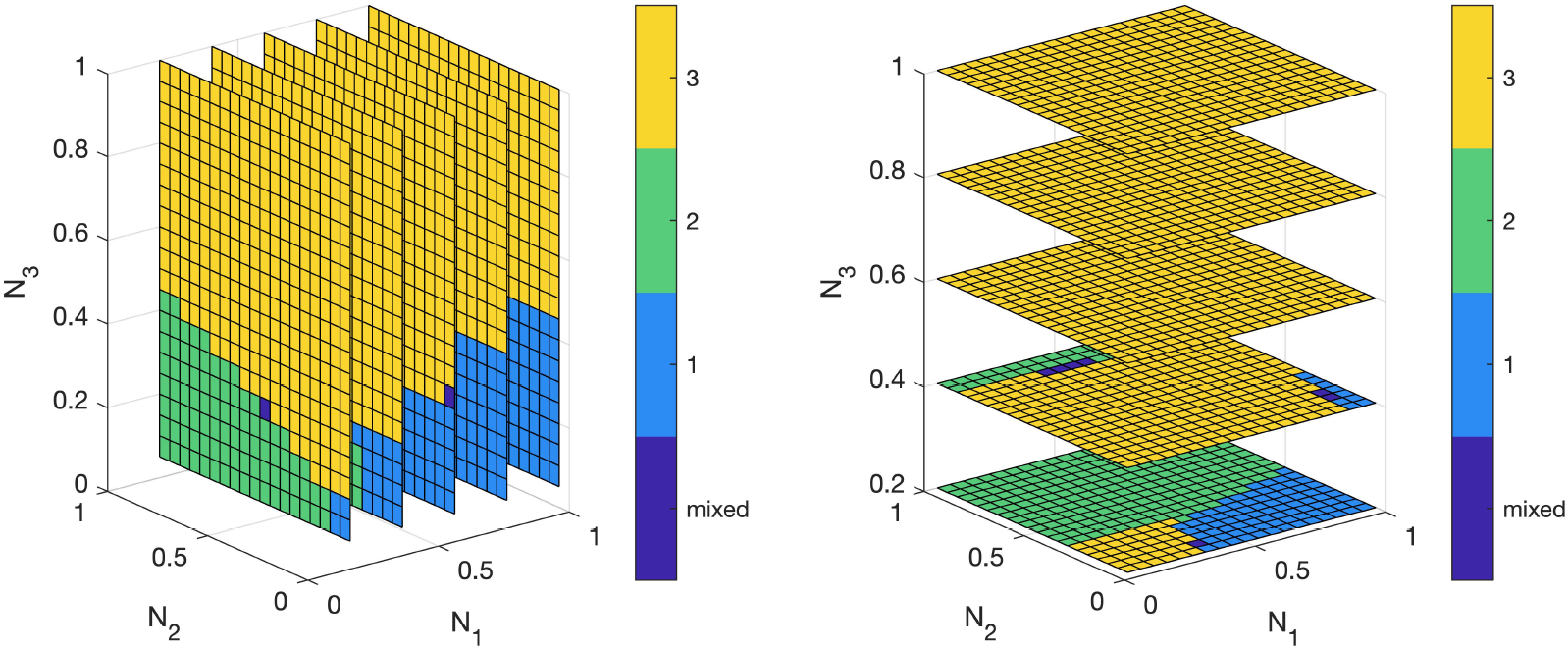
The long-term dynamics of the full life cycle map, with model ingredients set to the *k* = 3 analogue of (19). The figures show the winning cohort (1,2 or 3), or whether a mixed steady state was reached. Mixed steady states appear on the boundaries of the basins of attraction for each year class, and involve the year classes on either side of these boundaries. Note that SYC dynamics still predominates.

The derivative at the trivial fixed point *N*_3_ = 0 corresponds to multiplication by

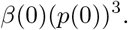

If this quantity exceeds 1, the population cannot go extinct, whereas there is an Allee effect when this quantity is less than one and yet part of the graph lies above the 45° line (see the green and orange curves in Figure 3 for examples).

Some possible choices for *p* and *β* include

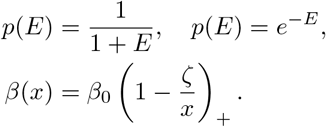

This choice for *β* is strictly increasing only where it takes strictly positive values and equals zero in a neighbourhood of *x* = 0, so there is certainly an Allee effect. It corresponds to the total population size of adults being reduced at a constant rate during a fixed time window. Alternatively, we can solve, with *P*, the predator density, as a parameter,

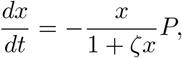

for a fixed period of time. This leads to an implicitly defined function *β*. For bifurcation diagrams of Single Year Class Maps, see Chapter V of (Davydova 2004).

Concerning dissipativity (Hypothesis 1), let for 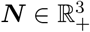

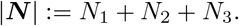

Then (15) implies that

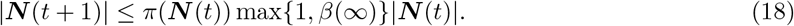

If for some choice of *R* > 0 and *ϵ* > 0 the inequality

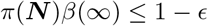

holds for all |***N***| ≥ *R*, then the set

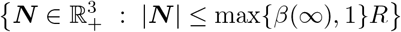

is forward invariant and attracts all orbits. Indeed, any orbit starting outside the set 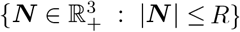 has to enter this set and once inside this set we can use (18) and the fact that *π*(***N***) ≤ 1 to conclude that, if *β*(∞) > 1, it may again leave the ball with radius *R* but not the ball with radius *β*(∞)*R*.

## 5 Discussion

The discrete time population dynamics of semelparous species with one reproduction opportunity per year, is described by a special kind of Leslie matrices, viz. those for which in the first (= reproduction) row only the last element is non-zero. This reflects that an individual born in a certain year reproduces, if at all, exactly *k* years later, where *k* is the length of the life cycle. So the population splits into *k* reproductively isolated year classes. Mathematically this shows up in the fact that the *k* times iterated matrix is diagonal, so fully reducible.

Although they are reproductively isolated, the year classes do interact by competition for resources. When resource availability is constant in time, each year class does as well or bad as any other year class. But when resource availability fluctuates in time, the phase of the life cycle relative to resource peaks and troughs can be decisive for success or failure. Thus one year class can drive other year classes to extinction by inducing, for instance, periodic environmental conditions. The work of Davydova and coworkers (Davydova 2004, Davydova et al. 2003; 2005, Diekmann et al. 2005) focused on this phenomenon.

The present paper is inspired by (Machta et al. 2019) and analyses a model such that, over a full life cycle, competition for food is neutral with respect to the year class distinction. But the model includes a second form of density dependence: it assumes that, due to predator satiation effects, per capita reproduction success of adults is larger when the cohort of adults is larger. So individuals belonging to a large cohort get much offspring while the negative impact of cohort size on survival probabilities is affecting individuals of all other year classes equally strongly, when measured over a full life cycle. Thus the ‘strongest’ year class becomes even stronger and eventually drives all other year classes to extinction and SYC (single year class) dynamics (Davydova et al. 2003, Mjølhus et al. 2005) gets established. The idea that a combination of predator satiation and intraspecific competition among larvae gives rise to single year class dynamics goes at least back to Hoppensteadt and Keller (1976) and Bulmer (1977), and has been demonstrated in many of the models in the literature by way of numerical experiments. The assumption of uniform competition introduced in (Machta et al. 2019) allows, as we have shown above, to actually prove that SYC dynamics results for large classes of initial conditions. In our opinion, the precise nature of the relationship between mechanisms and phenomena is better revealed by a proof than by simulations.

To explain, in the context of the model, that several species coexist in synchrony would probably require the assumption that both the functions *π* and the functions *β* for the various species are proportional. So this is still rather puzzling.

We now briefly consider some of the ecological evidence supporting the two chief modelling ingredients.

The long larval stages associated with periodical insects likely result from development constrained by certain abiotic factors such as low temperature, poor food availability, and large adult body size (Danks 1992, Hellövaara et al. 1994). *Magicicada* larvae feed underground on xylem found in plant roots. Xylem is a transport liquid in plants and is poor in nutrients. The competition for this shared food resource thus likely affects all cohorts feeding on them. Cicada nymphs have been shown to be uniformly spatially distributed, suggesting that cohorts do indeed compete for xylem (Williams and Simon 1995).

Predator satiation is well-documented for periodical cicadas (Williams and Simon 1995, and many references therein). The first cicadas to emerge still face a high predation pressure, with as much as 40% eaten within the first days. As numbers increase dramatically in the days following the start of the event, with up to 3,5 million adult cicadas per hectare, per capita predation pressure drops practically to zero (Williams et al. 1993). The predators, mainly birds such as cuckoos, woodpeckers, jays and other larger birds, do not increase much in number during the year of the outbreak, but several show increased population sizes in the one to three years to follow (Koenig and Liebhold 2005). It has also been shown that for several of these species, predator population sizes are in fact lower right before an outbreak event, thus decreasing predation pressure and allowing the insects to benefit even more from their numerical dominance (Koenig and Liebhold 2013).

In this paper we have focussed purely on the problem of elimination of year classes and how a single cohort arises from a starting situation with multiple cohorts. In our modelling framework we have not allowed for so-called stragglers, insects that have a longer developmental time and thus emerge in the year after an outbreak. This has been investigated recently using predominantly numerical simulations by Blackwood et al. (2018).

The enigma of periodical insect species is not confined to *Magicicada*, but is found also in several other insect orders, notably among *Xestia* moths, and in some well-known beetle species such as *Melonontha* cockchafer beetles (Hellövaara et al. 1994). Life spans may vary between species, and even between populations of the same species. *Magicicada* are exceptional, however, in their life spans, which are either 13 or 17 years, and are among the longest of all insect life spans.

There are several intriguing open theoretical questions regarding periodical insects. The adult insects, having developed under ground over a period of 13 or 17 years, all emerge from their burrows within an extremely short time span, usually between 7 and 10 days (Williams and Simon 1995, and references therein). Just how they synchronise so precisely remains unclear (Williams and Simon 1995), although temperature cues have been suggested.

In some outbreak years, as much as half of all individuals fail to develop into adults, and eclose the next year (White and Lloyd 1979). This has been associated with extremely poor nutrition conditions. Differences in the developmental rate of individual larvae are apparently common, with late-developing nymphs ‘catching up’ before emerging as adults with the rest. To incorporate such phenomena would require developing a physiologically structured population model, with developmental rate directly influenced by food availability and competition (de Roos and Persson 2013).

## Acknowledgments

Part of the work was done during the Mathematical Biology semester at the Mittag-Leffler Institute. We thank the organisers, in particular Torbjörn Lundh and Mats Gyllenberg, for making it such a success. We also thank two anonymous reviewers for their careful reading and suggestions to improve the manuscript.

## Appendix

For the class of models considered in (Machta et al. 2019), it is shown in that paper that a steady state, with more than one year class present, is necessarily unstable. The aim of this appendix is to show, by way of an example, that for the class of models considered here it is, in contrast, possible to have a stable steady state with multiple year classes. In order that simple computations suffice to reach this conclusion, we choose

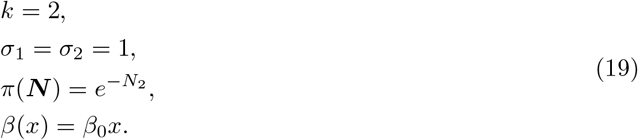

So we focus our attention on

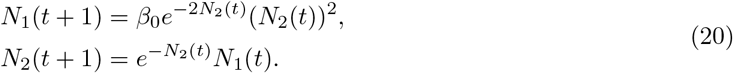

If 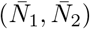 is a steady state, we should have

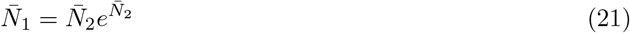

and

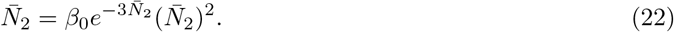

The trivial steady state (0,0) is locally stable but, due to positive density dependence incorporated in *β*, this does not exclude that nontrivial steady states exist. The equation

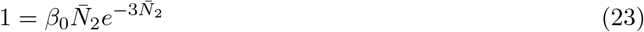

has for

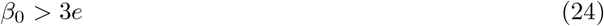

two positive solutions, one with 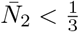 and one with 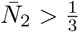. Each of these yields, when combined with (21), a nontrivial steady state. The corresponding linearized system is given by

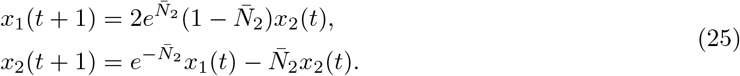

The Jacobi matrix

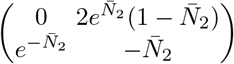

has trace 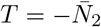 and determinant 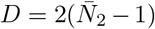. Since *T* < 0, the stability conditions are *D* < 1 and *T* + *D* + 1 > 0 and amount to

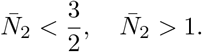

So if we think in terms of bifurcations that happen when the parameter *β*_0_ is increased, the scenario is as follows:

a. at *β*_0_ = 3*e* a saddle-node bifurcation of nontrivual steady states happens, but both steady states are unstable
b. the larger, with respect to both components, of the two steady states undergoes a period-doubling bifurcation at *β*_0_ = *e*^3^; for 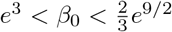 this steady state is stable.
c. at 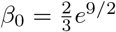 this steady state loses stability in a Neimark-Sacker bifurcation.

For completeness, let us have a brief look at SYC dynamics. If *N*_1_(*t*) =0 we obtain

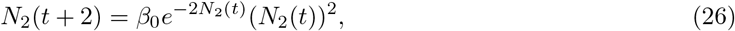

with stable trivial steady state and two nontrivial steady states for *β*_0_ > 2*e* arising by saddle-node bifurcation at *β*_0_ = 2*e* with value 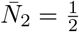. The linearized recursion is

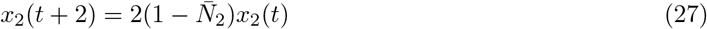

so we see that the larger of the two is stable for

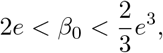

losing its stability by period doubling at the upper boundary of this window.

It appears that the bifurcation diagrams of the “all year class” steady state and of the “single year class” steady state are in no way related to each other.

It is easy to repeat the bifurcation analysis with *π* in (19) replaced by

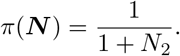

It turns out that in this case the “two year class” steady state is unstable for all parameter values.

In Figure 4 we visualize the outcome of numerical experiments with *k* = 3 and model ingredients such that two year classes can coexist in a stable fixed point of the full life cycle map. By symmetry there are three such fixed points. It appears that the domains of attraction of these three fixed points are rather small. As a rule, there is an ultimate winner.

